# Predicted Impact of the Viral Mutational Landscape on the Cytotoxic Response against SARS-CoV-2

**DOI:** 10.1101/2021.07.04.451040

**Authors:** Anna Foix, Daniel López, Michael J. McConnell, Antonio J. Martín-Galiano

## Abstract

The massive assessment of immune evasion due to viral mutations that potentially increase COVID-19 susceptibility can be computationally facilitated. The adaptive cytotoxic T response is critical during primary infection and the generation of long-term protection. Potential epitopes in the SARS-CoV-2 proteome were predicted for 2,915 human alleles of 71 HLA class I families. Allele families showed extreme differences in number of recognized epitopes, underscoring genetic variability of protective capacity between humans. Up to 1,222 epitopes were associated with any of the twelve supertypes, that is, allele clusters covering 90% population. Among them, the B27 supertype showed the lowest number of epitopes. Epitope escape mutations identified in ~118,000 NCBI isolates mainly involved non-conservative substitutions at the second and C-terminal position of the ligand core, or total ligand removal by large recurrent deletions. Escape mutations affected 47% of supertype epitopes, which in 21% of cases concerned isolates from two or more sub-continental areas. Some of these changes were coupled, but never surpassed 15% evaded epitopes for the same supertype in the same isolate, except for B27, which reached up to 33%. In contrast to most supertypes, eight particular allele families mostly contained alleles with few SARS-CoV-2 ligands. Isolates harboring cytotoxic escape mutations for these families co-existed geographically within sub-Saharan and Asian populations enriched in these alleles. Collectively, these data indicate that independent escape mutation events have already occurred for half of HLA class I supertype epitopes. However, it is presently unlikely that, overall, it poses a threat to the global population. In contrast, single and double mutations for susceptible alleles may be associated with viral selective pressure and alarming local outbreaks. This study highlights the automated integration of genomic, geographical and immunoinformatic information for surveillance of SARS-CoV-2 variants potentially affecting the population as a whole, as well as minority subpopulations.

**AUTHOR SUMMARY:** The cytotoxic T response, a type of immune response dependent upon an individual's genetics that does not require antibodies, is critical for neutralizing SARS-CoV-2 infection. The potential bypass of the cytotoxic T response by mutations acquired by the virus after one year of the pandemic is therefore of maximal concern. We have approached the complexity of human variability and more than 100.000 viral genomes in this respect using a computational strategy. We have detected numerous mutations in these genomes that mask some viral regions involved in the cytotoxic response. However, the accumulation of these changes in independent isolates is still too low to threaten the global human population. In contrast, our protocol has identified mutations that may be relevant for specific populations and minorities with cytotoxic genetic backgrounds susceptible to SARS-CoV-2 infection. Some viral variants co-existed in the same country with these human communities which warrants deeper surveillance in these cases to prevent local outbreaks. Our study support the integration of massive data of different natures in the surveillance of viral pandemics.

## Introduction

Mutations in the severe acute respiratory syndrome coronavirus 2 (SARS-CoV-2) leading to increased susceptibility are of extreme concern. Given the slow pace of vaccination in some geographic regions, enhanced primary infection by strains that evade immune detection might worsen the significant health and socioeconomic burden caused by the COVID-19 pandemic.

Long term protection from viral infection relies on a competent adaptive response. Adaptive protection includes the coordinated activation and memory of three adaptive response compartments. These branches consist of the humoral response, driven by antibodies synthesized by B cells, and the two types of cellular response, driven by CD8^+^ and CD4^+^ lymphocytes that recognize viral peptides bound to human leukocyte antigen (HLA) class I and II molecules, respectively (1). Antibodies neutralize the virus, for example, by specific binding to the spike protein and inhibit binding to the ACE2 receptor expressed in the lung (2). CD8^+^, or cytotoxic, T lymphocytes directly kill SARS-CoV-2 infected cells through the secretion of pore-forming proteases and the induction of programmed cell death (3). Finally, CD4^+^, T helper lymphocytes, play a pivotal stimulatory role for both antibody-led and cytotoxic activities. An effective cellular response is associated with prompt and efficient protection during primary and successive SARS infections (4–6). Moreover, the cellular and humoral responses are long-lasting (7) and elicit immunoprotective memory (8,9).

Naïve cytotoxic lymphocytes are stimulated through the presentation of specific proteolyzed fragments of antigens, or epitopes, bound to HLA class I molecules in the membrane of infected cells. Some peptides of approximately nine residues are generated by cleavage of intracellular pathogen proteins and bound in the endoplasmic reticulum to the apical antigenic groove of the monomeric chain of HLA class I. Once in the membrane, these ligands can be recognized by the T-cell receptor of T CD8^+^ lymphocytes that start the maturation process. After activation and division of a sufficient subsets of mature CD8^+^ T cells, the subject may be protected against the particular virus as SARS-CoV-2 at a cellular level (3).

Severe COVID-19 outcomes have been associated with aging and co-morbidities such as hypertension and diabetes mellitus (10). However, how host genetic factors influence the disease is still largely unknown. In this respect, the adaptive cellular response is strongly influenced by host genetics. Notably, HLA class I genes are among the most multiple and variable genes in humans. The HLA class I system consists of three loci for which over 17,000 alleles have been reported (11). These alleles are further grouped into phylogenetic families and, some of them, into supertypes that shared comparable ligands (12). Overall, this huge allelic diversity provides the human species with an enormous capacity to detect different antigens from virtually any pathogen.

HLA class I epitope pool screenings with SARS-CoV-2 sequences have been carried out for specific countries (13–15). This kind of experimental information is stored in repositories like the Immune Epitope Database and Analysis Resource (IEDB) (16), which allows for the global analysis of potential mutation-evasion events. Certain HLA alleles have been associated with permissiveness to SARS-CoV-2 infection, such as B*44 and C*01 families in Italy (17), and B*15:27 and C*07:29 in China (18). However, these data still do not evenly represent genetic differences in susceptibility at the global population level. Since analyzing thousands of HLA alleles is experimentally unrealistic (19), the confines of the human SARS-CoV-2 cytotoxic ligandome can be explored by *bona fide* computation approaches in a neutral manner. For instance, Nguyen *et al*. identified HLA-B*46:01 as the less efficient allele for presenting SARS-CoV-2 epitopes among 145 alleles by using an immunoinformatic approach (20).

Widespread infection with SARS-CoV-2 at the global level provides the virus with great opportunity to explore the mutational space. Some of these changes may be selected based on immunological evasive advantages. In this respect, how the genetic variability of the virus can affect individuals carrying different HLA class I alleles is currently unknown. Viral mutations that dramatically decrease binding affinity of epitopes to HLA class I molecules can act as escape mutants and alter the cellular immunity, with important implications for clinical evolution of the infection (21). How many and which mutations an isolate must acquire in order to evade the adaptive cytotoxic responses of the general population remains an open question. Such emerging capacity to bypass the cytotoxic response would presumably not follow a categorical binary pattern but a gradient dependant on underlying individual genetics.

The goal of this study is two-fold. First, we have interrogated how SARS-CoV-2 mutations can affect predicted HLA class I binding at the global population level, and second, how existing mutations influence the response of specific sub-populations that harbor alleles with few SARS-CoV-2 epitopes. For that, we have taken advantage of the strength of computational methods to design a protocol that generates and operates on formatted data. This allowed us to conduct an exhaustive analysis that involved over 2,900 human HLA class I alleles and ~118,000 viral genomes. The knowledge acquired here may help to understand the current status of the human cytotoxic defense in the context of the pandemic and to promptly identify emerging strains that require close monitoring.

## Results

### Predicted vs validated SARS-CoV-2 HLA class I epitopes

To obtain a more complete insight of the cellular cytotoxic response to SARS-CoV-2, HLA class I epitopes from the SARS-CoV-2 reference proteome were predicted by the universal netMHCpan 4.1 EL algorithm (22). These included all medium-strong peptide binders for 2,915 human alleles grouped into 71 families of the HLA-A (21 families, 886 alleles), HLA-B (36 families, 1412 alleles), HLA-C (14 families, 617 alleles) loci available in this software. The predicted full SARS-CoV-2 ligandome for HLA class I reached 5,224 independent epitopes.

Data complexity reduction by clustering alleles into families can cause some information loss. However, the degree of intra-family coherence, that is, the percentage of matching epitopes between two alleles of the same family with respect to the total predicted by both alleles, reached 61.3 ± 19.2% (mean ± SD). In contrast, the inter-family coherence, that is, the average matching epitopes after all-against-all family comparison), was only 3.0 ± 1.6%. This supports that, despite the existence of intra-family differences, the allele family cluster stratum is acceptable for a global view of HLA epitopes.

Families showed a drastic difference in the number of predicted epitopes (Fig. 1A). Globally, families of A and C loci showed higher values than B loci families. In particular, A*01, A*23, A*24, C*12 and C*14 families surpassed 300 epitopes on average, whereas B*46, B*82 and B*83 were below five. Twenty-six alleles, several from the B*46 family, were not associated with any predicted epitope. These computational predictions are in line with antecedent observations concerning great differences between HLA class I alleles in the response to the SARS-CoV-2 reference strain (18,20). Some families were linked to exclusive epitope pools but others shared overlapping SARS-CoV-2 ligandomes (Fig. 1B).

**Fig 1.**
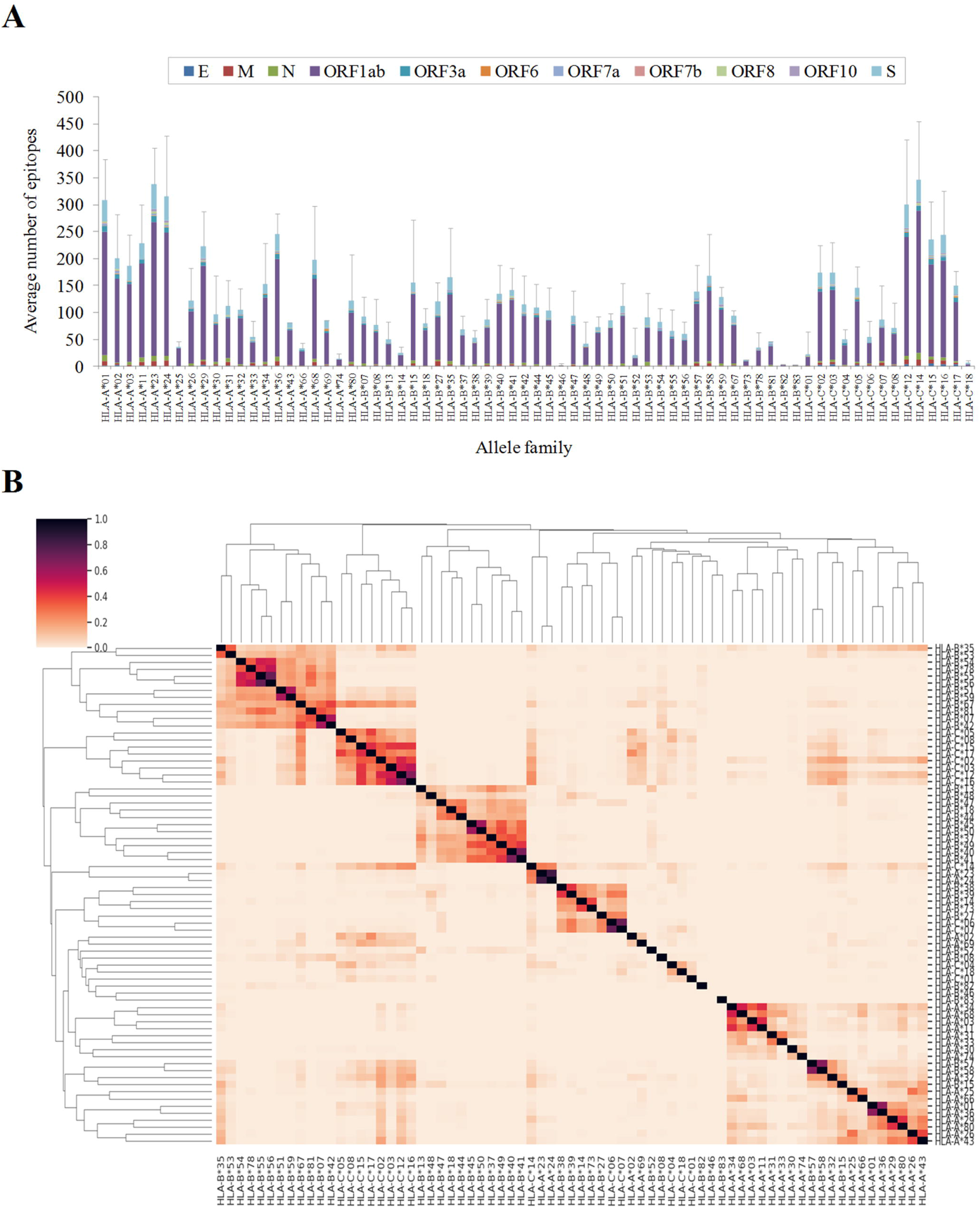
Number and degree of overlap between SARS-CoV-2 epitopes for different HLA-class I allelic families. **(A)** Average number of predicted HLA class I epitopes by allele family and protein. The standard deviation resulting from all proteins is indicated as a single error bar. **(B)** Hierarchical clustering and associated heatmap indicating the degree of inter-family epitope correlation. Color intensity expresses the Jaccard index for the epitope intersection between all family pairs. Perfect location match between epitopes calculated by netMHCIpan 4.1 EL with score ≥ 0.5 and rank ≤ 0.5 were utilized to calculate intersection and union. Intra-family conserved epitopes (≥ 50% alleles in the family by exact match) are in Supplementary Table S1.

All viral proteins theoretically generated HLA class I epitopes. On average, 1.19 epitopes per allele family (those identified for ≥ 50% alleles in the family) and 100 residues were identified in the SARS-CoV-2 proteome. Among polypeptides with ≥ 75 residues, the M protein carried higher (1.41 epitopes per family and 100 residues) and N lower (0.71) epitope densities, respectively.

Epitope predictions were compared to 760 experimentally validated 8-12mer epitopes for HLA class I included in the IEDB dataset. Up to 90% of validated epitopes perfectly matched predicted epitopes, for at least one allele, at the stringent thresholds applied. There was high correlation (r^2^ = 0.87, polynomic fit) between the number of predicted and validated epitopes for the allele (Fig. 2A). However, several alleles from the A*02 family were comparatively over-represented while B*27 and B*39 alleles were under-represented in the validated dataset. Differences for sequence and number of validated ligand datasets were evident between whole families (Fig. 2B). Globally, the ratio between validated and predicted epitopes was significantly higher for families of the A locus (0.35 ± 0.12) with respect to those of the B (0.31 ± 0.13, *P* < 0.001 Student's t-test) and the C (0.29 ± 0.10, *P* < 0.001) loci. Forty-four alleles from 13 families did not show any validated experimental epitope, a higher number than allele and families without predicted epitopes. Despite invaluable studies that contributed data with relatively large and distributed datasets (23), experimental screenings may be slightly biased by the low frequencies of some alleles in the cohorts analyzed. Overall, computational and experimental approaches may be complementary and beneficial for the global characterization of the SARS-CoV-2 cytotoxic response.

**Fig 2.**
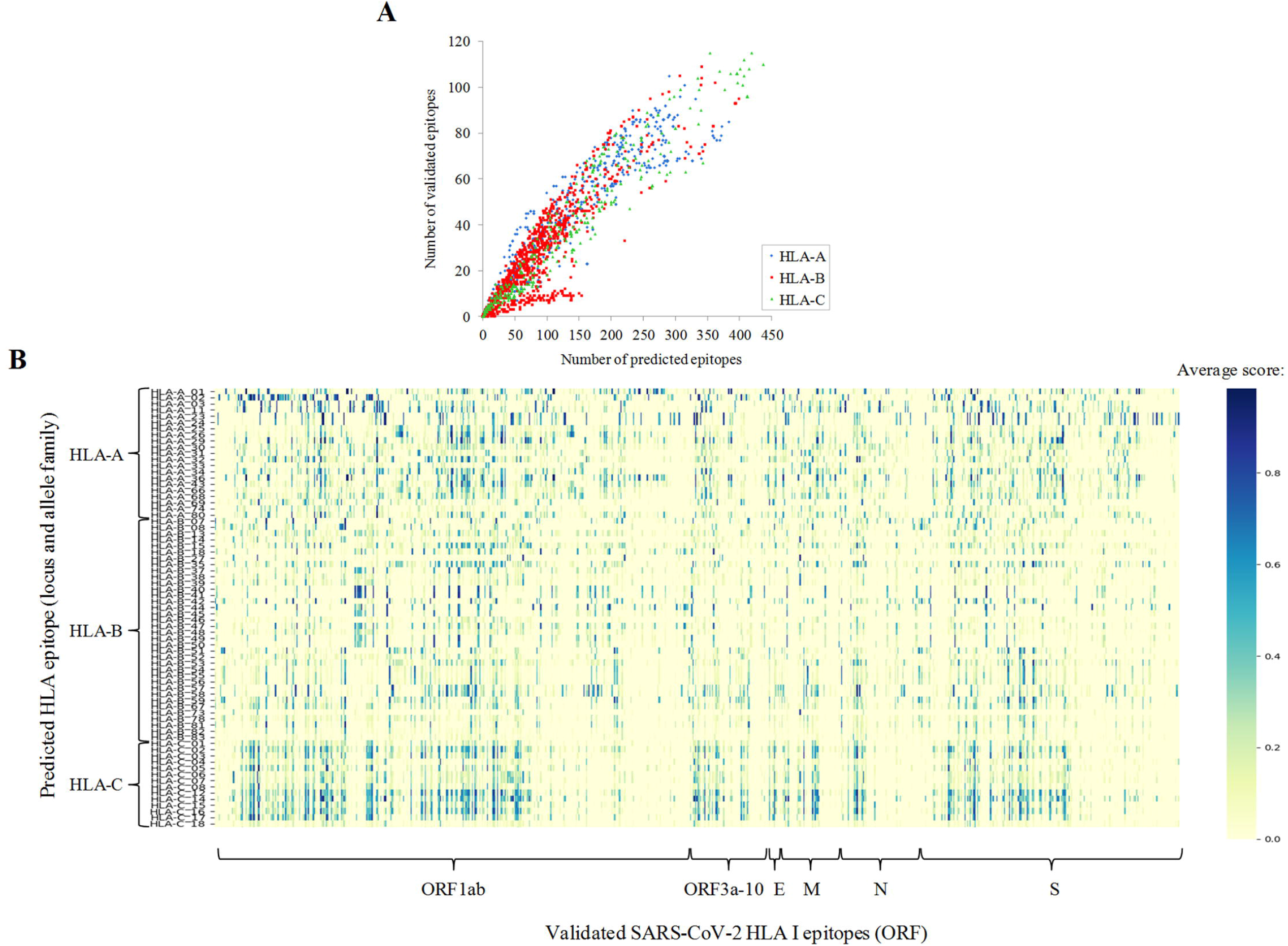
Comparison between predicted and validated epitopes. **(A)** Number of predicted epitopes (score ≥ 0.5 and rank ≤ 0.5) versus validated epitopes per allele. **(B)** Heatmap showing the family average score (any score, rank ≤ 2) for validated HLA class I epitopes. Predicted epitopes with perfect matches with validated epitopes stored in the IEDB are indicated in Supplementary Table S1.

### Supertypes show very different number of SARS-CoV-2 supermotifs

Alleles, from the same or different families, that bind similar epitopes are functionally grouped into twelve, so-called, supertypes (12). Supertypes cover >90% of the world population regardless of ethnicity. In our dataset, 1,222 (23.4%) of all non-redundant epitopes were able to cover ≥ 50% of the alleles associated with at least one supertype, i.e. are supermotifs (24) (Fig. 3A, Supplementary Table S2). Moreover, twenty supermotifs covered three or more supertypes (Table 1). On average, 11.1 supermotifs were identified per 100 residues of the viral proteome. Among ORFs with ≥ 75 residues, the M (13.5 supermotifs per 100 residues) and the N (5.0 supermotifs per 100 residues) proteins showed the highest and the lowest concentration of supermotifs, respectively. The number of supertypes was unevenly represented since, for instance, the “A01 A24” and “A24” supertypes were associated with >250 supermotifs, while others as “A01” or “B07” with around 50 supermotifs, and the “B27” supertype showed only 12 (Fig. 3B).

**Fig 3.**
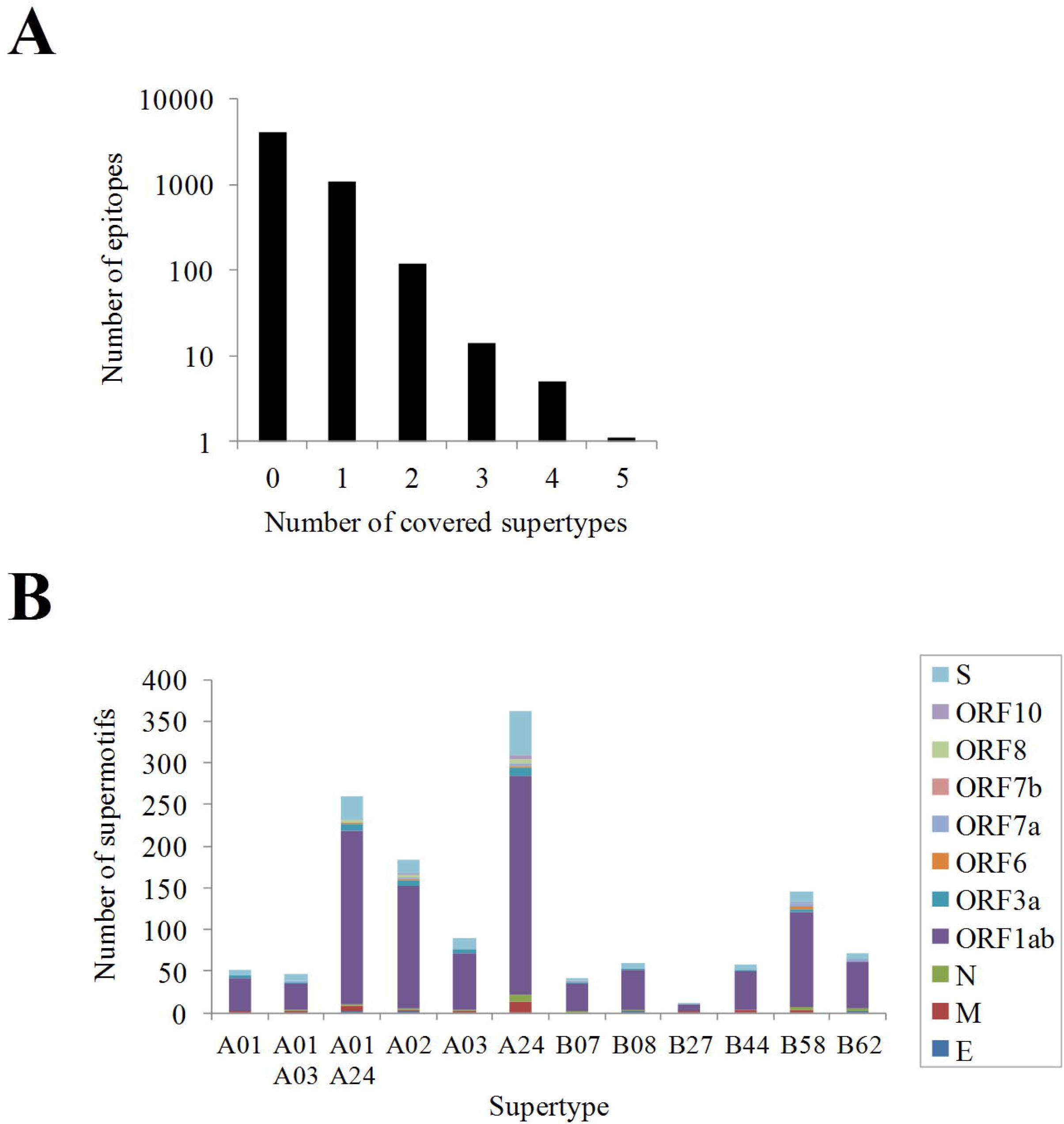
SARS-CoV-2 supermotifs. **(A)** Distribution of supermotifs according to the number of supertypes covered. **(B)** Number of supermotifs per supertype detailed by protein antigen.

**Table 1.**
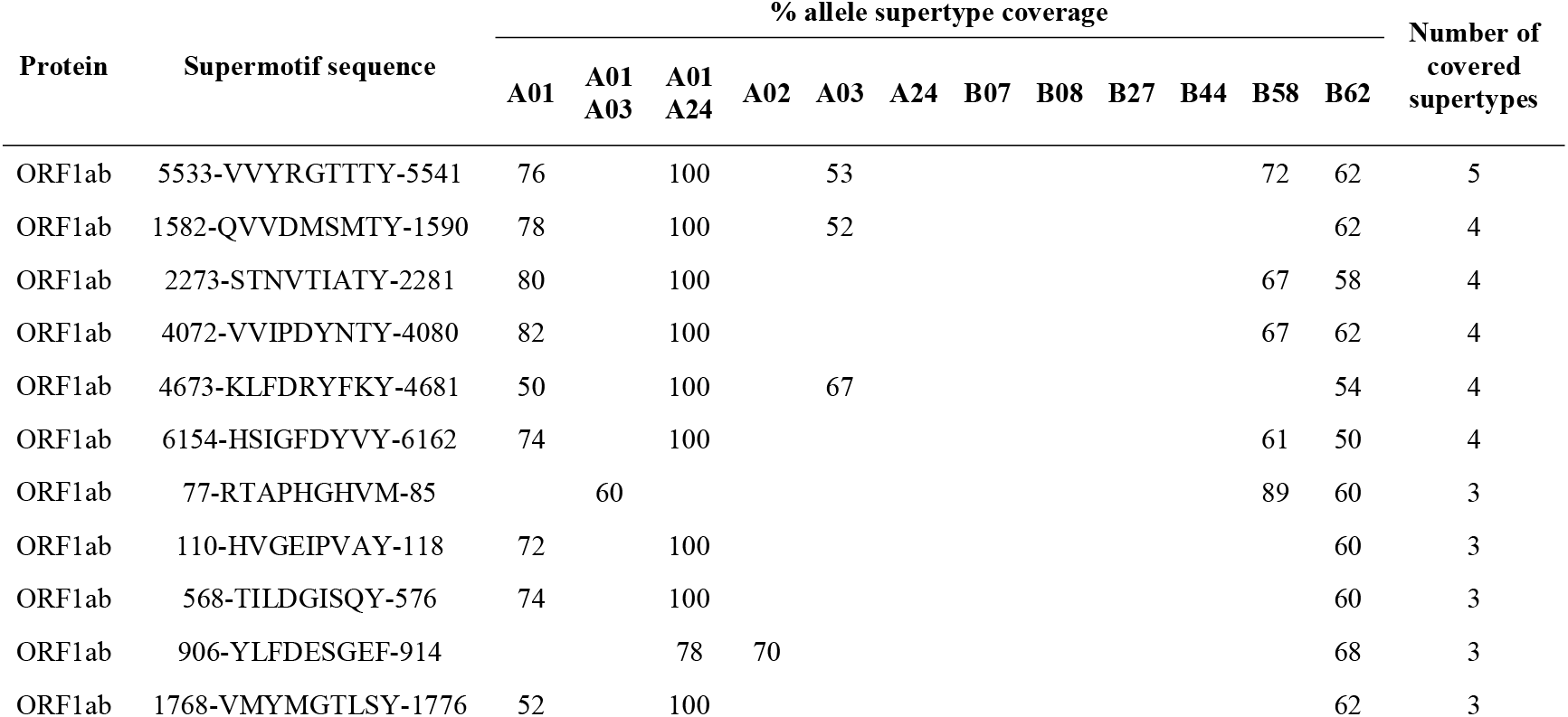
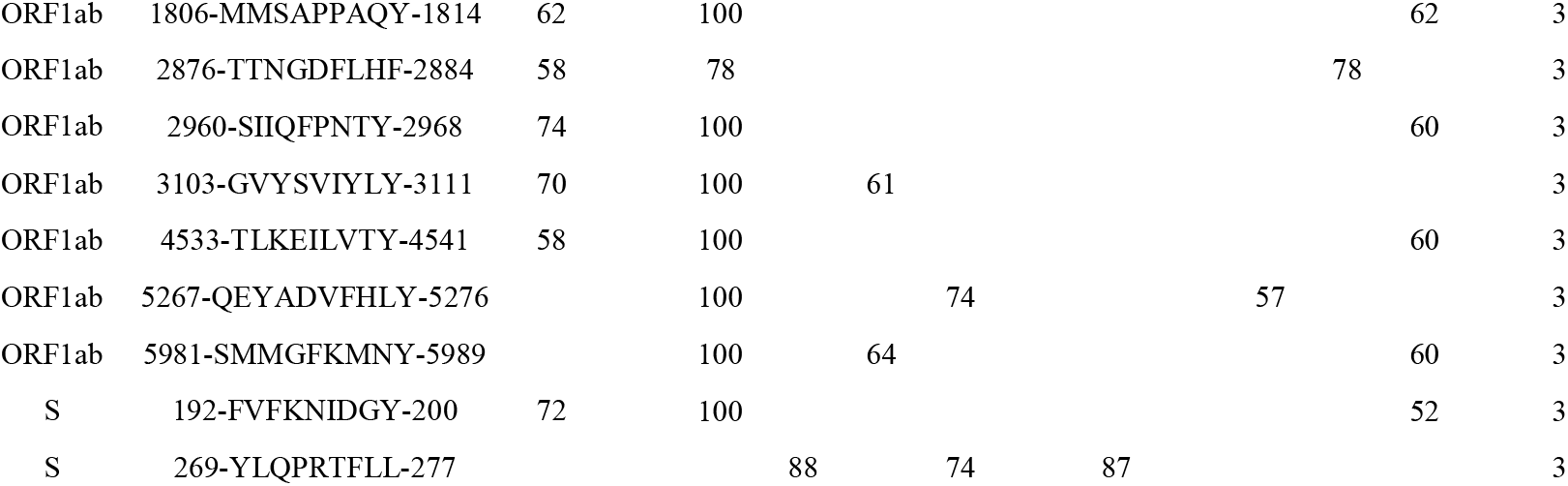
SARS-CoV-2 HLA class I supermotifs involving three or more supertypes.

### Recurrent mutations can affect HLA class I epitopes

Calculations performed up to this point have included only the original Wu-han-1 reference strain. However, viruses are continuously evolving entities where HLA class I ligand recognition can be dynamically subjected to extensive mutation-selection processes. Key mutations could produce cytotoxic escape variants by reducing affinity or even deleting HLA class I ligands and, then, influencing the ability of CD8+ lymphocytes to clear the infection (25). To assess this possibility, mutations were identified in 117,811 SARS-CoV-2 isolates from 87 countries covering 21 out of the 22 sub-continental areas of the United Nations M49 geoscheme (https://unstats.un.org/unsd/methodology/m49/). A total of 1,128,631 genetic alterations with respect to the Wu-han-1 reference strain were identified. These involved 28,512 unique residue substitutions in 9,723 positions. A total of 78% of unique substitutions were non-conservative, those concerning distinct amino acid classes (Fig. 4A, left). Up to 26,231 deletions of 1 to 193 residues and 127 insertions, ranging between 1 and 8 residues, were also detected (Fig. 4B). Substitutions were much more prevalent than deletions (Fig. 4C). In contrast, deletions affected a higher number of epitopes (Fig. 4C). Nevertheless, the degeneration of ligand binding is expected to counteract many substitutions and point deletions affecting epitopes, leading to a negligible effect on the cytotoxic response.

**Fig 4.**
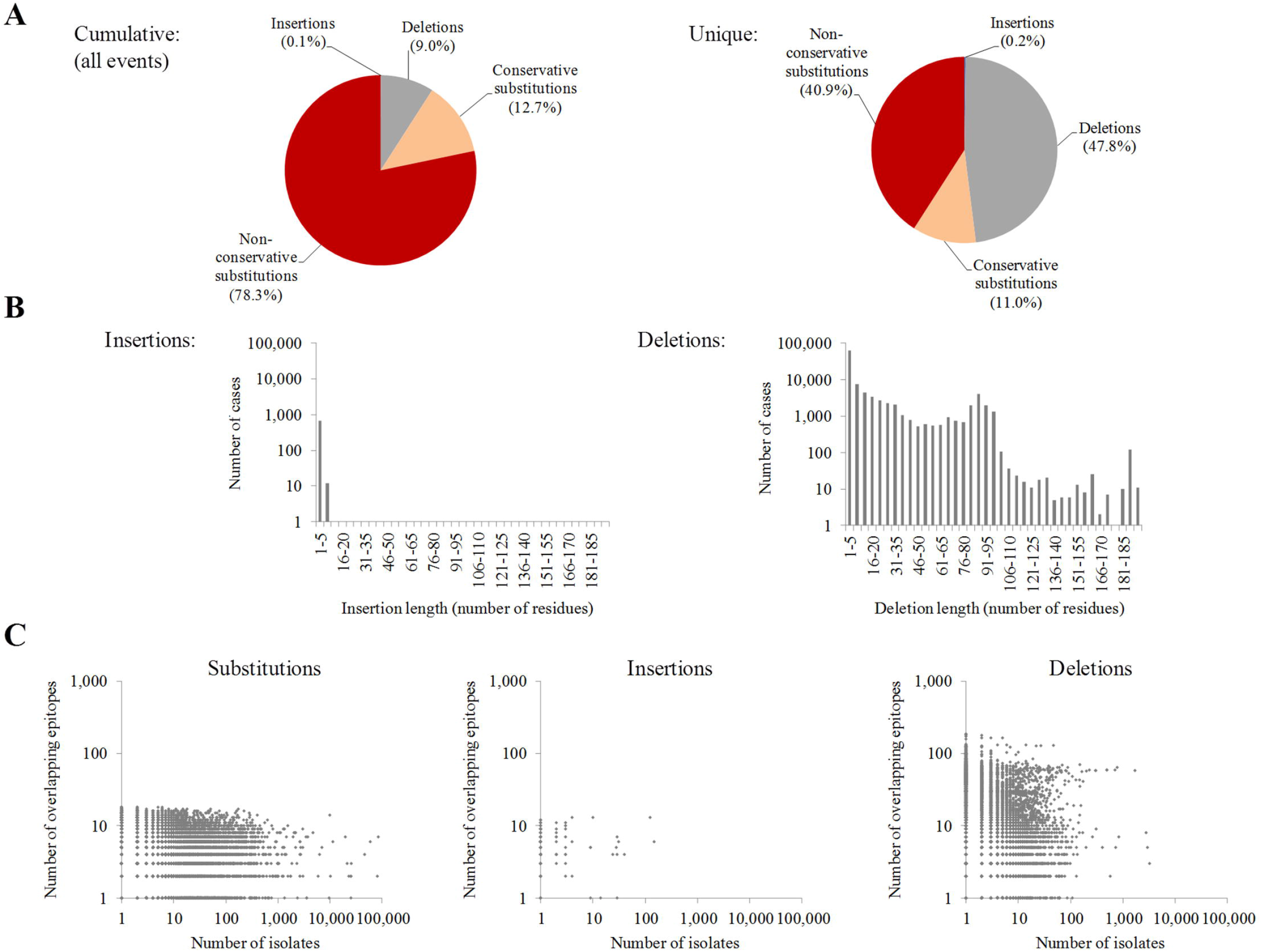
Global mutation analysis in NCBI SARS-CoV-2 genomes. **(A)** Proportion of cumulative and unique residue mutation events in SARS-CoV-2. **(B)** Length distribution of insertions (left) and deletions (right). **(C)** Number of isolates and number of epitopes which location overlap to substitutions (left), insertions (center) and deletions (right).

### There are mutations for most supermotifs but only a fraction causes epitope escape and are geographically distributed

All 1,224 supermotifs carried some type of mutation in least one isolate. Nevertheless, a central question is to what extend these changes have high impact in the context of the cytotoxic response of the worldwide population.

Mutations were scrutinized using three criteria: recurrence, binding affinity reduction and geographical dissemination of isolates carrying them. For this, a series of incremental selective criteria on all the genetic changes observed was applied: (Filter 1) presence of the mutation in ≥ 2 isolates if the mutation was a substitution, or ≥ 5 isolates if the mutation was an insertion or a deletion, since these are more be resultant of sequencing errors; (Filter 2) drastic reduction of supermotif binding. Changes in the second (P2) and C-terminal (P9 in core nonamers) positions preferentially perturb binding affinity, in particular when these changes involve amino acids of different physicochemical classes (non-conservative) (Fig 5A). This can be explained by residues in these positions intimately interact with the selective B and F pockets in the groove of the HLA molecule. However, the epitope disturbing capacity of mutations in these positions is not an exact rule (Fig. 5A). Thus, the actual impact on binding was explicitly recalculated in the mutated sequence and quantified using recommended thresholds (see Materials and Methods); and (Filter 3) detection of the mutation in isolates from different M49 world regions.

**Fig 5.**
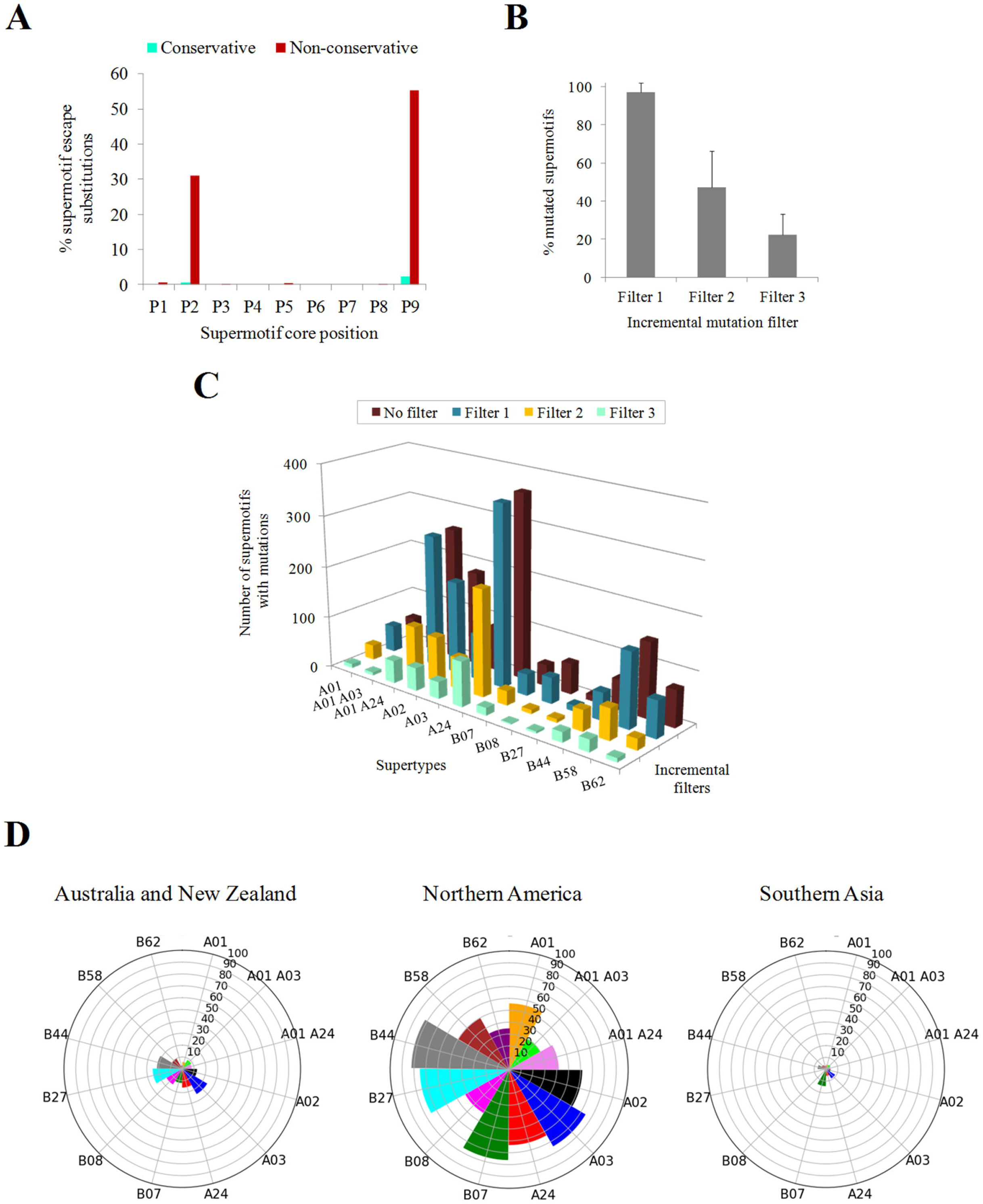
Supermotif escape mutations. **(A)** Influence of supermotif core position and residue conservation in the epitope escape capacity of substitutions. **(B)** Average percentage of escape supermotifs by any mutation type after incremental filter application. **(C)** Absolute number of mutated supermotifs for each supertype after incremental filter application. **(D)** Nightingale rose charts indicating the percentage of escape supermotifs in prevalent M49 zones. Only mutations involving ≥2 isolates in the M49 were considered. Only M49 zones with ≥ 5% escape supermotifs for at least one supertype are shown.

Expectedly, the fraction of escape supermotifs substantially decreased as more selective criteria were applied (Fig. 5B). Only 22.1% of supermotifs contained mutations that satisfied all the stringent criteria, that is, show recurrent mutations that cancel the HLA class I binding and are found in isolates over several sub-continental zones. Such high-impact changes affected differently to various supertypes. The A03 supertype was more affected with 36.7% while only 3.3% of B08 supertype supermotifs showed escape and disseminated mutations (Fig. 5C, Table S2). The “Australia and New Zealand” and, in particular, “Northern America” M49 zones presented isolates with mutations in a substantial part of supermotifs (Fig. 5D).

Recurrent mutations may have great relevance if they affect more than one supertype. Thirty substitutions that disabled binding affinity of universal supermotifs appeared in 70 or more isolates (Table 2, Supplementary Table S3). Among them, was the W152C change in the spike protein in 3,455 USA isolates, which removed four supermotifs of two supertypes. The W6974V (ORF1ab) substitution found in 211 isolates of four M49 areas, destroyed three supermotifs of two supertypes. Notably, the P5828 and K6980 (ORF1ab) positions showed two different recurrent escape mutations each.

**Table 2.**
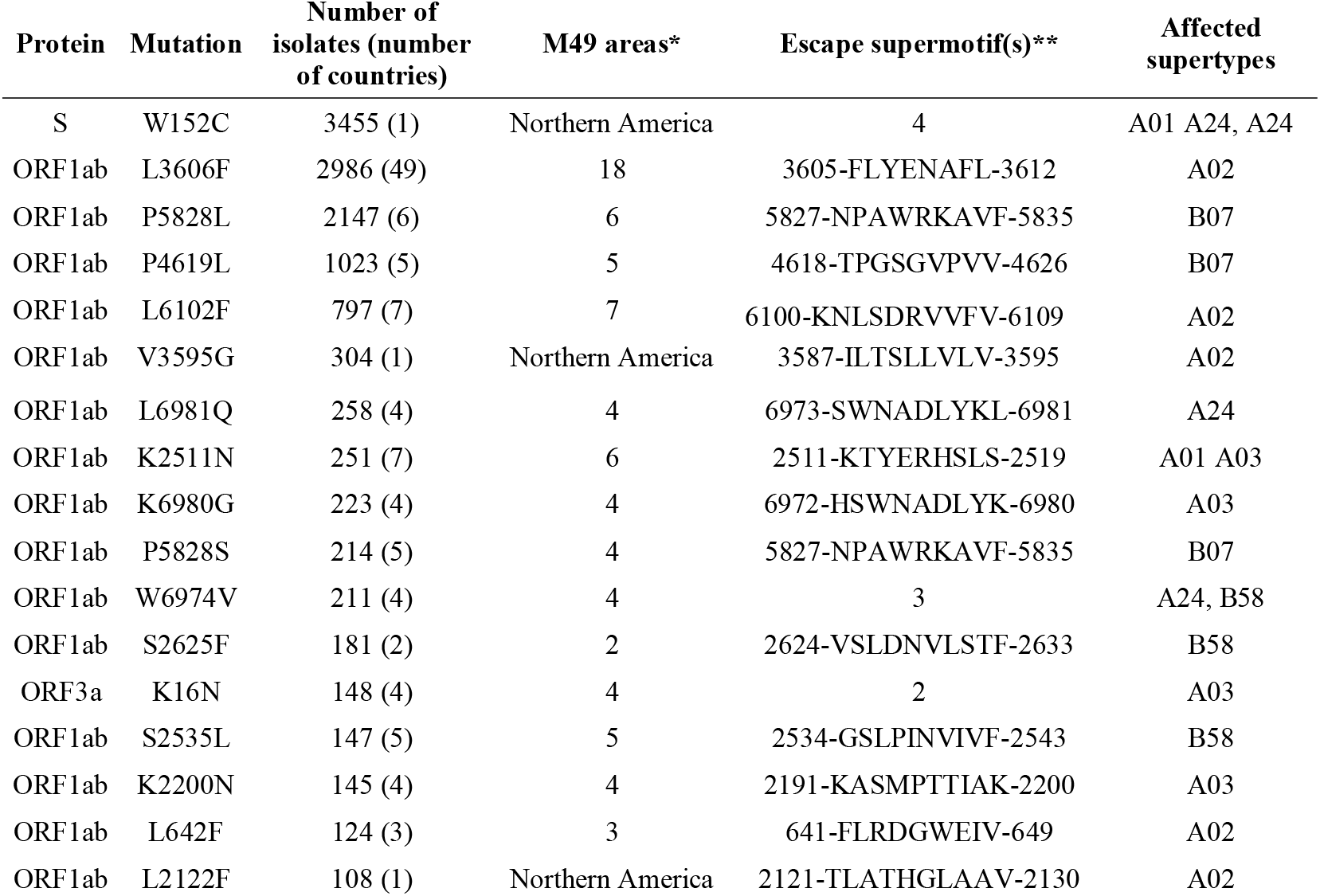
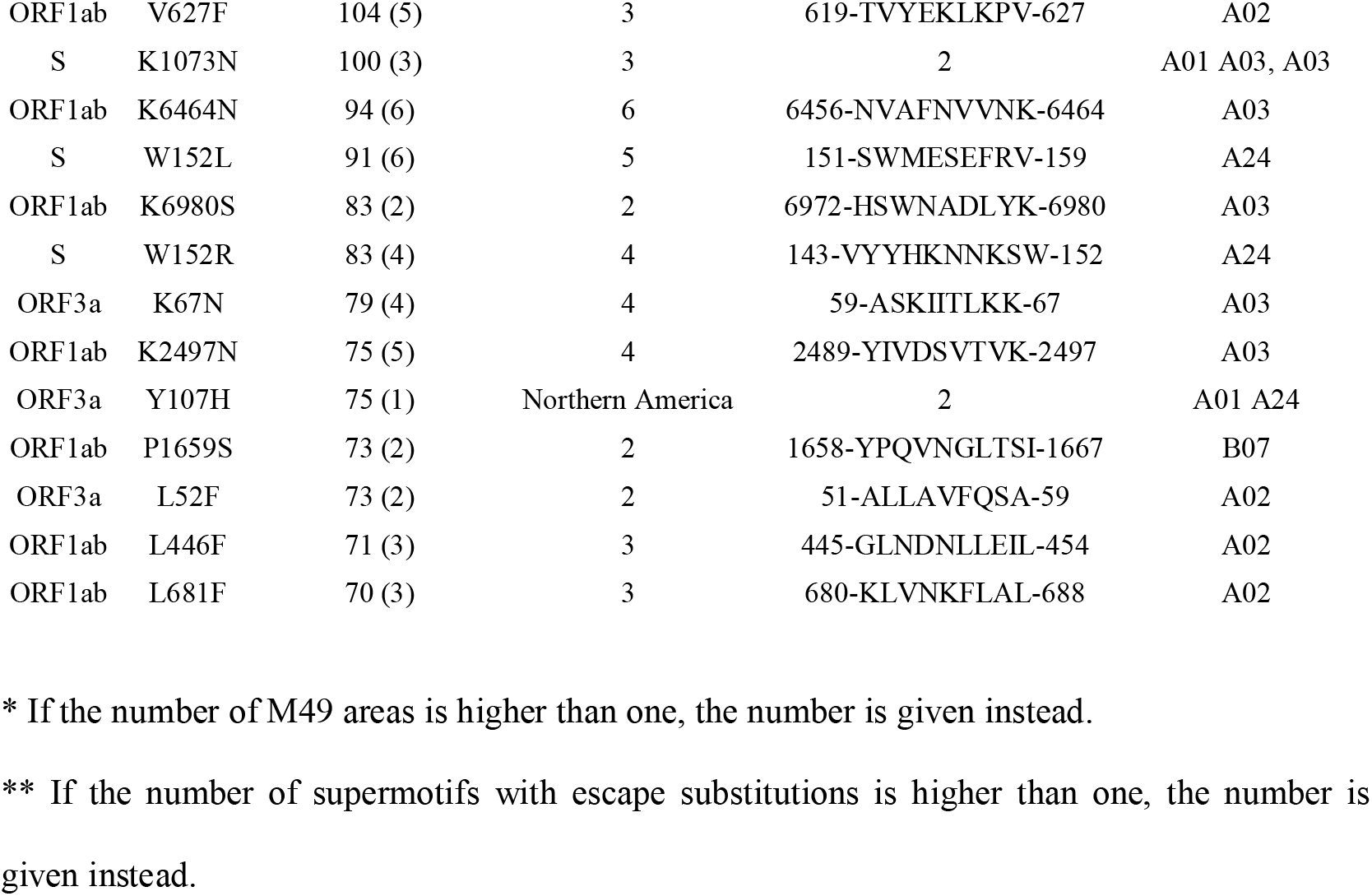
Top-30 recurrent supermotif escape substitutions.

Comparatively, insertions played an almost global negligible effect. Only 6965::SKF and 6981::KEGQ (ORF1ab) decreased the HLA binding, affected one supermotif and supertype each, and were in more than 10 isolates (Table 3).

**Table 3.**
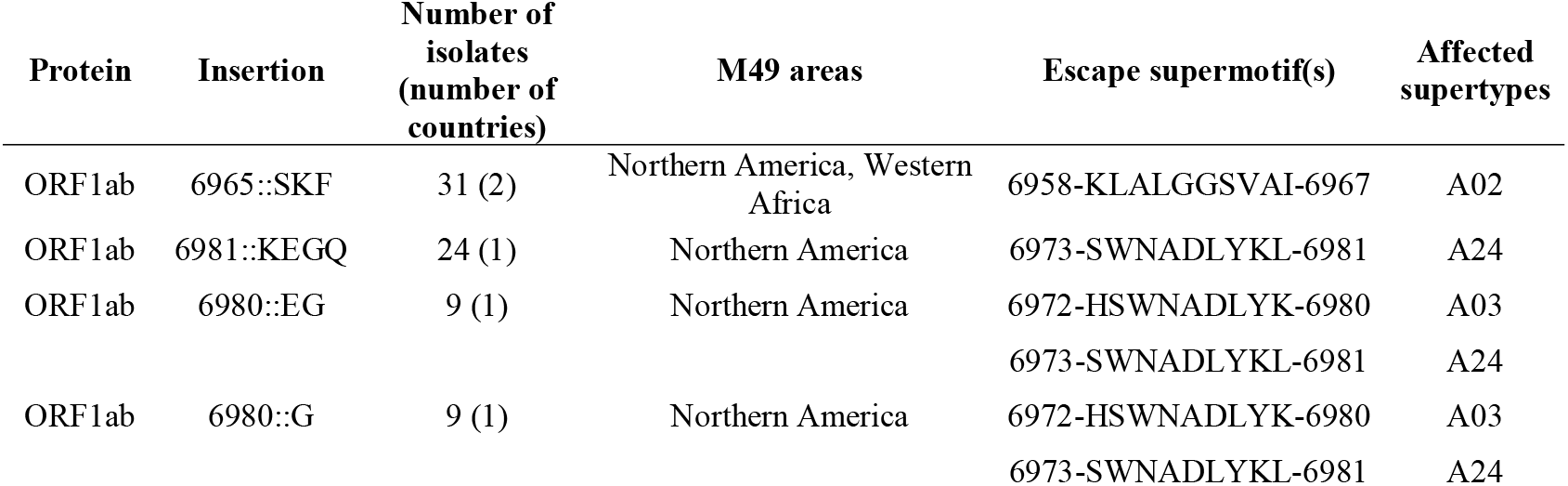
Recurrent supermotif escape insertions.

| Protein | Insertion | Number of isolates (number of countries) | M49 areas | Escape supermotif(s) | Affected supertypes |
| --- | --- | --- | --- | --- | --- |
| ORF1ab | 6965::SKF | 31 (2) | Northern America, Western Africa | 6958-KLALGGSVAI-6967 | A02 |
| ORF1ab | 6981::KEGQ | 24 (1) | Northern America | 6973-SWNADLYKL-6981 | A24 |
| ORF1ab | 6980::EG | 9 (1) | Northern America | 6972-HSWNADLYK-6980 | A03 |
|  |  |  |  | 6973-SWNADLYKL-6981 | A24 |
| ORF1ab | 6980::G | 9 (1) | Northern America | 6972-HSWNADLYK-6980 | A03 |
|  |  |  |  | 6973-SWNADLYKL-6981 | A24 |

Deletions showed a binary pattern (Table 4 and Supplementary Table S4). On the one hand, short deletions (≤ 3 residues) affected many proteome zones and mostly concerned a single supermotif. For instance, the predominant Δ145 (S protein), Δ363 (N protein) and Δ141-143 (ORF1ab) deletions were in this group. On the other hand, long recurrent deletions (>80 residues) removed up to twelve supermotifs from seven supertypes and tended to occur in discrete proteome hotspots, namely the 2791-2883 (28-120 of nsp4) and the 6338-6436 (413-511 of nsp14A2_ExonN) regions of ORF1ab.

**Table 4.**
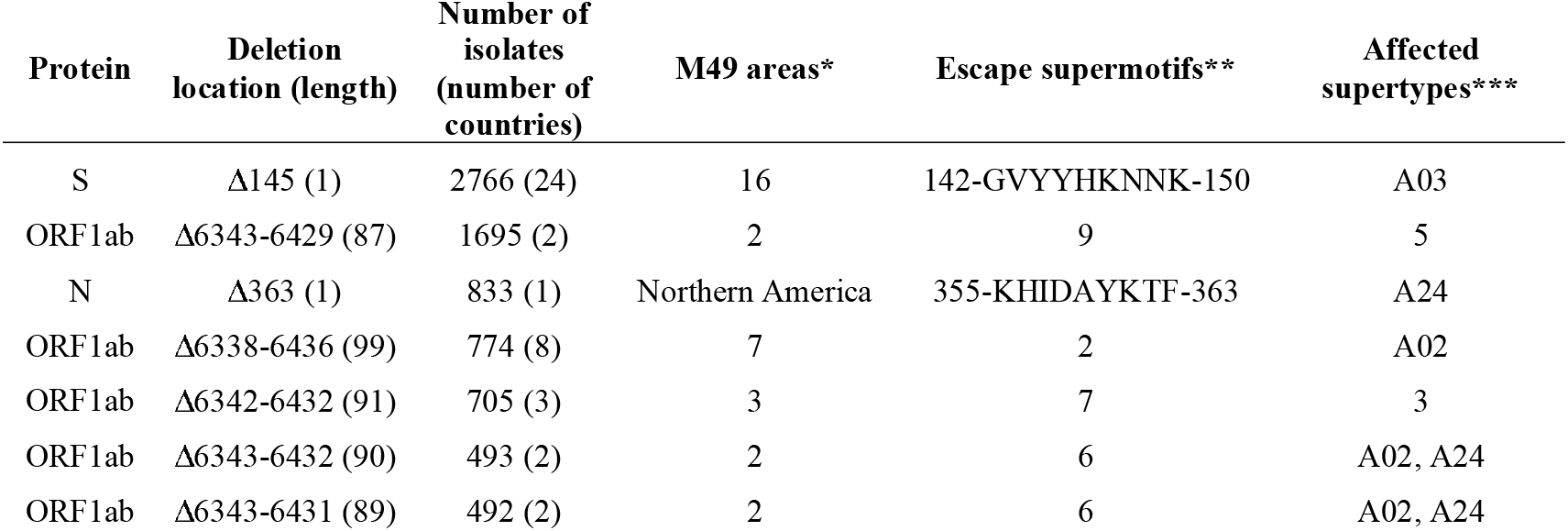
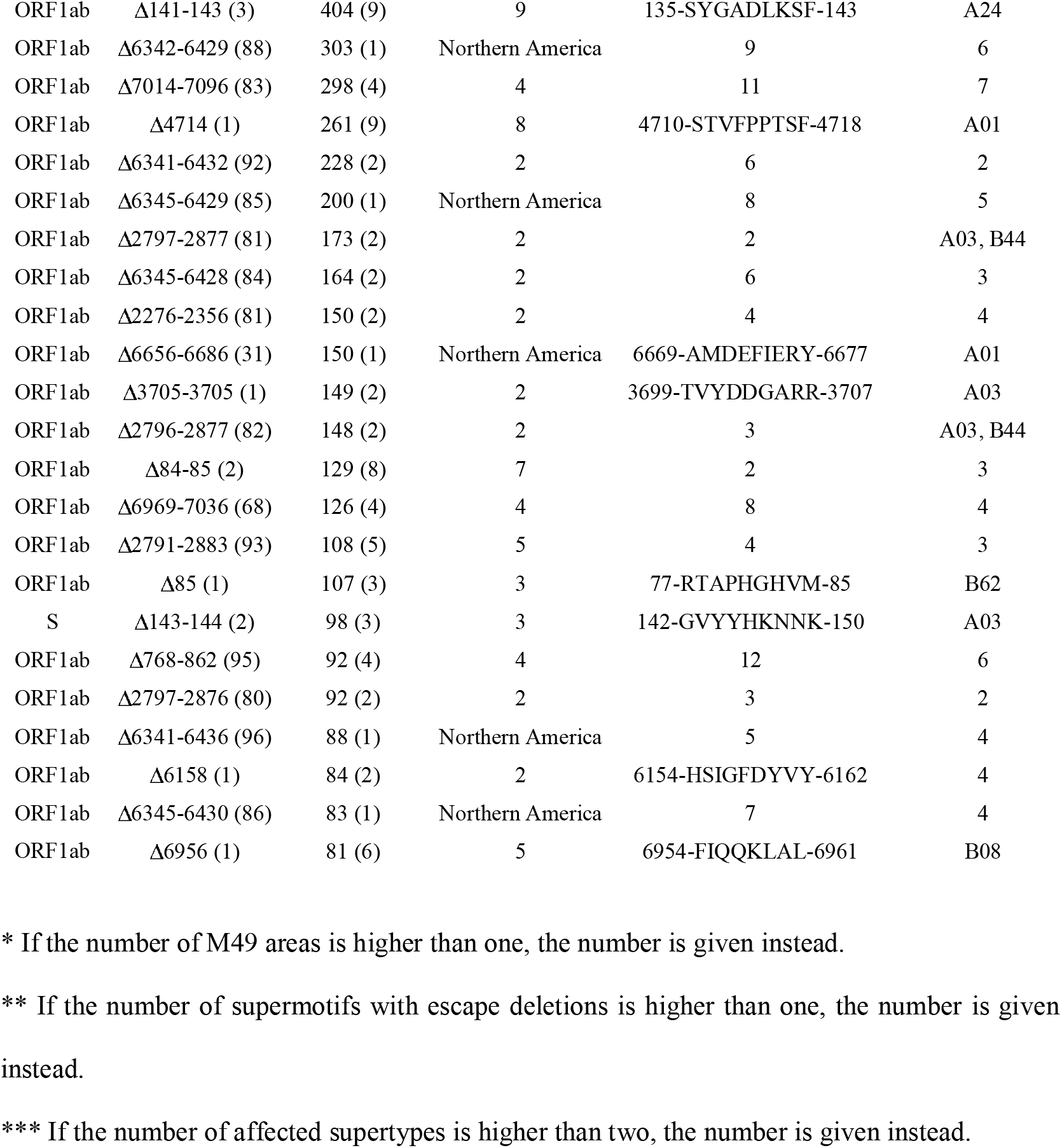
Top-30 recurrent supermotif escape deletions.

### Only a few supermotif escape mutations coexist in the same isolate

Beyond prevalence, these mutations may show distinct combinatorial preferences for simultaneously co-occurring in the same isolates. This information was utilized to detect nine independent mutation networks of 2-44 mutations. Several supermotif mutations were linked through a few spread mutations acting as hubs: W152C (S protein), L3606F (ORF1ab, L37 in nsp6_TM) and four long recurrent deletions in the 6342-6432 range (ORF1ab, nsp14A2_ExonN protein)(Fig. 6).

**Fig 6.**
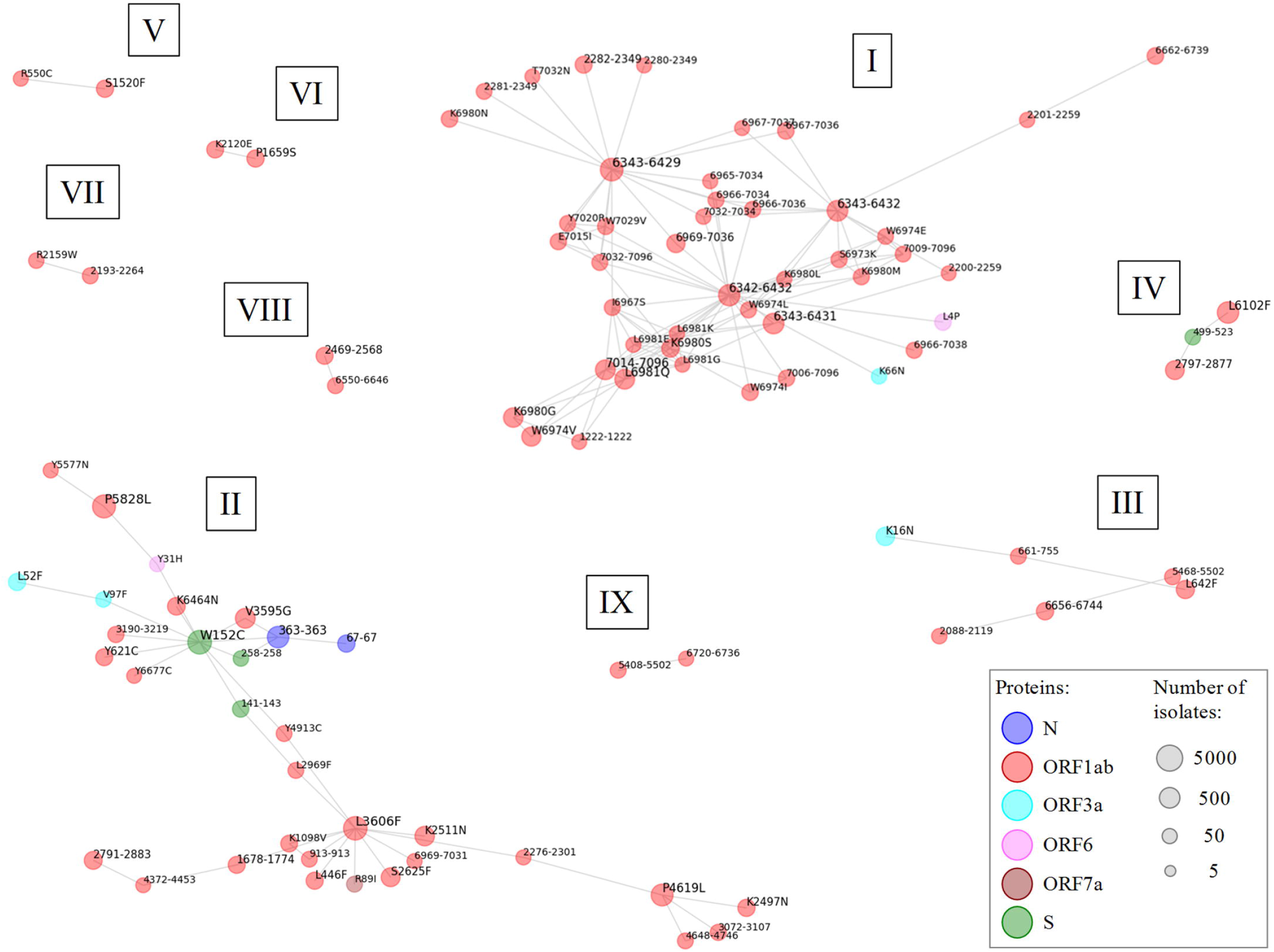
Networks of coupled supermotif escape mutations. Undirected unweighted graphs showing coupled supermotif escape mutations. Sub-networks are named with roman numbers. Nodes correspond to mutations that were substitutions (position and residue change) or deletions (residue range). No coupled insertions were detected. The node color indicates the antigen protein. The sphere diameter reflects the amount of isolates harboring the mutation. Nodes represent mutations carried by ≥ 25 isolates. Edges represent co-existence of a mutation pair in ≥ 20% isolates of all those carrying at least one of the mutations.

Mutated lineages were also analyzed at the isolate level. Ultimately, isolates enriched in supermotif escape mutations may evade immune system response and disseminate quickly. There was a direct relationship between the number of substitutions in an isolate and the number of supertype alleles altered, which may be mostly attributed to neutral RNA replication errors. Importantly, 7347 isolates conveyed mutations in ≥ 5 supermotifs (Table 5 and Supplementary Table S5) and in 1027 cases affected ≥ 5 supertypes.

**Table 5.** Top isolates showing supermotif escape mutations.

| Accession | Total number of mutations | Number of escape supermotifs | Number of affected superotypes | Ratio escape supermotifs per mutation | Collection date | Country |
| --- | --- | --- | --- | --- | --- | --- |
| MW673525 | 40 | 19 | 5 | 0.475 | 08/02/2021 | USA |
| MT577016 | 297 | 18 | 7 | 0.061 | 2020 | India |
| MW694016 | 31 | 15 | 9 | 0.484 | 11/02/2021 | USA |
| MT451283 | 276 | 14 | 8 | 0.051 | 24/03/2020 | Australia |
| MW156473 | 51 | 14 | 7 | 0.275 | 28/07/2020 | Australia |
| MT451279 | 142 | 13 | 8 | 0.092 | 24/03/2020 | Australia |
| MT451436 | 134 | 13 | 8 | 0.097 | 26/03/2020 | Australia |
| MW689154 | 62 | 13 | 7 | 0.210 | 13/02/2021 | USA |
| MW653643 | 21 | 13 | 6 | 0.619 | 07/12/2020 | USA |
| MW525102 | 27 | 13 | 4 | 0.481 | 10/01/2021 | USA |
| MW406716 | 58 | 12 | 9 | 0.207 | 24/06/2020 | USA |
| MW190139 | 49 | 12 | 8 | 0.245 | 16/07/2020 | USA |
| MW228176 | 68 | 12 | 8 | 0.176 | 16/06/2020 | USA |
| MW406699 | 62 | 12 | 8 | 0.194 | 24/06/2020 | USA |
| MW542158 | 75 | 12 | 8 | 0.160 | 13/01/2021 | USA |
| MW725850 | 27 | 12 | 8 | 0.444 | 24/02/2021 | USA |
| MW449384 | 63 | 12 | 7 | 0.190 | 30/11/2020 | USA |
| MW474268 | 54 | 12 | 7 | 0.222 | 12/11/2020 | USA |
| MW617514 | 62 | 12 | 7 | 0.194 | 02/02/2021 | USA |
| MW704295 | 50 | 12 | 7 | 0.240 | 19/01/2021 | Bahrain |
| MW715548 | 71 | 12 | 7 | 0.169 | 19/02/2021 | USA |
| MW673420 | 38 | 12 | 6 | 0.316 | 09/02/2021 | USA |
| MW741583 | 26 | 12 | 6 | 0.462 | 23/02/2021 | USA |
| MW751588 | 30 | 12 | 6 | 0.400 | 04/03/2021 | USA |
| MW783199 | 68 | 12 | 6 | 0.176 | 02/03/2021 | USA |
| MW518131 | 34 | 12 | 5 | 0.353 | 03/01/2021 | USA |
| MW693032 | 18 | 12 | 5 | 0.667 | 05/11/2020 | USA |
| MW586153 | 16 | 12 | 4 | 0.750 | 28/01/2021 | USA |
| MW596067 | 32 | 12 | 4 | 0.375 | 29/01/2021 | USA |
| MW673042 | 33 | 12 | 3 | 0.364 | 07/02/2021 | USA |

The origin of most supertype-mutated isolates were USA and Australia (Fig. 7A). Among emergent isolates, a strongly mutated isolate (Assembly database entry: “MT577016", 297 mutations) from India stood out with 18 escape supermotifs corresponding to 7 supertypes. Notably, 16.4% of isolates, mostly showing only moderate mutational profiles (≥ 5 substitutions), presented ≥ 0.5 negated supermotifs per mutation. This feature suggests potential pressure for cytotoxic evasion by precise supermotif mutation in a subpopulation of isolates in the later cases. For instance, the MW586153 isolate collected in USA:SC showed 16 substitutions where twelve of them removed supermotifs from four supertypes. Other remarkable cases were three isolates (MW702787, MW702788 and MW702806) from the same county in USA:CA that showed eleven escape supermotifs from seven supertypes with only fourteen substitutions, suggesting an incipient evasive lineage.

**Fig 7.**
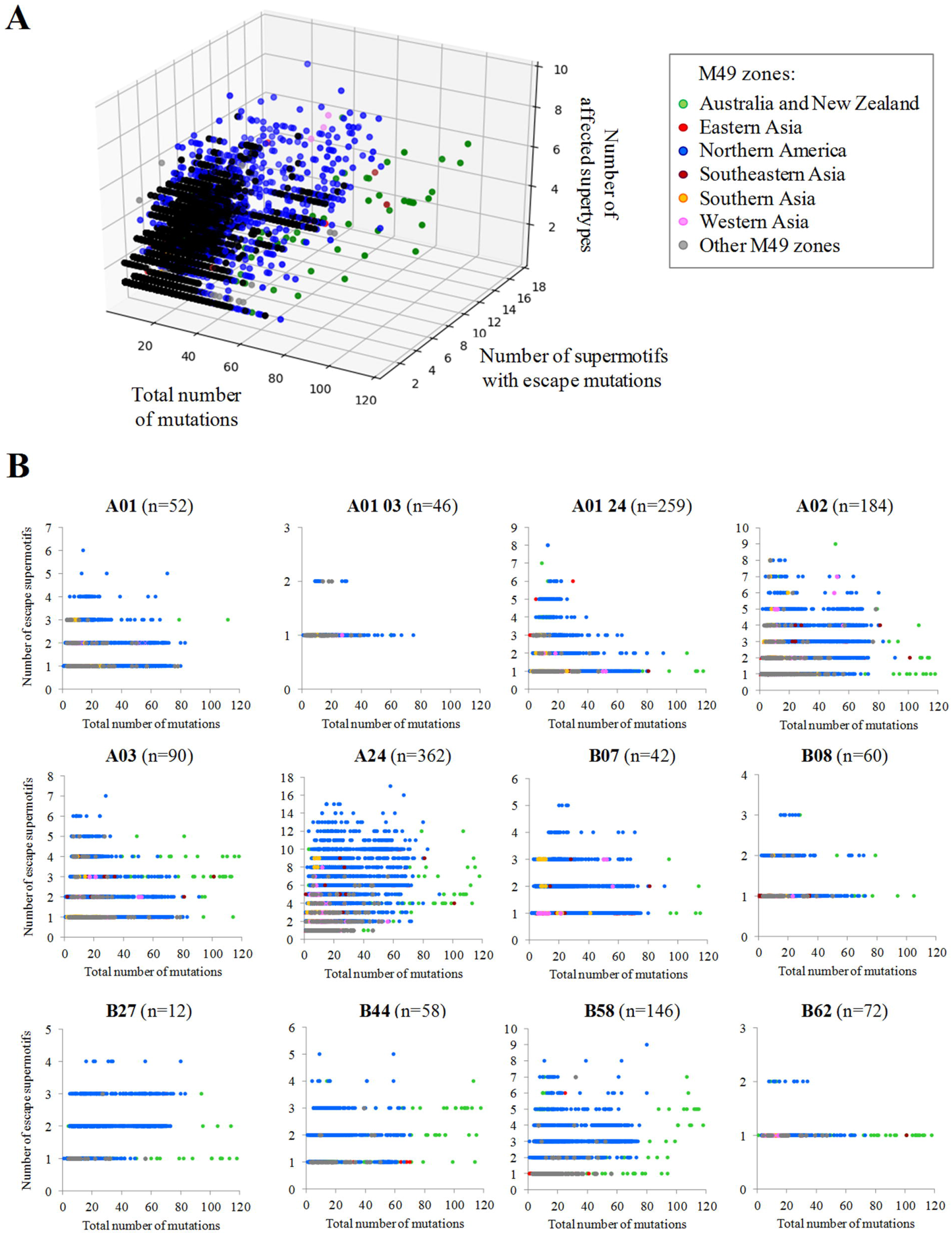
Isolates carrying different combinations of escape mutations. **(A)** Each point represents an isolate plotted according to the total number of mutations, the number of escape supermotifs and number of affected supertypes. Only isolates harboring three or more escape supermotifs are represented. **(B)** Chart panel indicating mutated isolates according to the number of escape supermotifs for each supertype. Isolates are colored by M49 zone of collection.

However, no isolates carried escape mutations for >15% supermotifs of specific supertypes (Fig. 7B). The only exceptions were three USA isolates with <25 mutations (including deletions) that invalidated four out of the twelve B*27 supermotifs. Seventeen USA isolates with ≤ 20 substitutions and no indels invalidated up to three B*27 supermotifs.

### Epitope escape mutations in families with scarce SARS-CoV-2 ligandomes

In contrast to most supertypes, some alleles did not shown affinity to any SARS-CoV-2 peptide or showed scarce SARS-CoV-2 peptide repertoires. In this light, 246 alleles (8.4%) of the three loci (HLA-A: 39 alleles; HLA-B: 143 alleles; HLA-C: 64 alleles) were predicted to bind with high affinity to twenty or less epitopes. These alleles belonged to 48 families which showed three possible patterns depending on their alleles with few SARS-CoV-2 epitopes was either the norm or the exception (Fig. 8A). Firstly, eight families contained ≥81% of alleles with few predicted epitopes and ≤ 22 epitopes per allele on average, and were deemed poor SARS-CoV-2-repertoire families. These families were A*74, B*46, B*52, B*73, B*82, B*83, C*01 and C*18. Remarkably, the combination of alleles of inefficient families for the three loci, the A*74:02-B*46:01-C*01:02 haplotype, has been detected with a 0.02% frequency in a Hong Kong sample. Secondly, and in contrast, most families analyzed contained only <17% alleles linked to few epitopes and ≥ 34 epitopes per allele on average. However, three of these families (B*08, B*15 and C*07) were large families that included ≥ 10 alleles with limited SARS-CoV-2 epitope sets. Finally, just two families behaved in a hybrid manner: B*14 (18% alleles with few epitopes; 23.7 epitopes/allele) and B*78 (43% alleles with few epitopes, 31.1 epitopes/allele).

**Fig 8.**
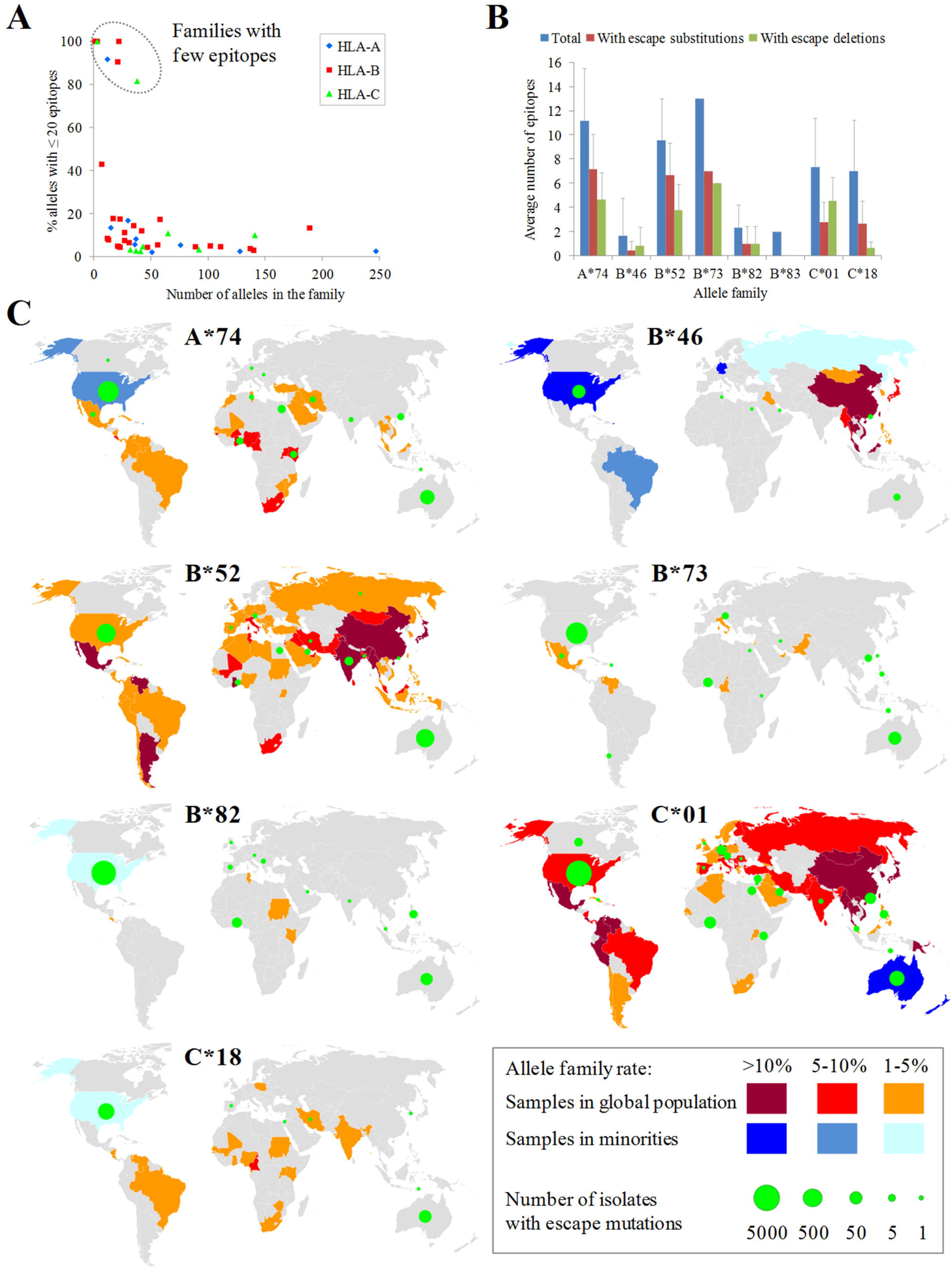
Escape mutations in allele families with fewest epitopes. **(A)** Number of alleles with ≤ 20 epitopes versus the total number of alleles for HLA families of the three loci. Families without any allele with ≤ 20 epitopes are not represented. **(B)** Average number of escape epitopes, either by substitutions or deletions, respect to the average total number of epitopes for the eight allele families with the fewest epitopes. **(C)** World map panel indicating the presence of population samples carrying alleles of the eight families with fewest epitopes and isolates with escape mutations for these families. Family allele frequencies are color ranked for both the majority population (red scale) and sub-population (blue scale) samples. Only the highest frequency sample per country was considered. B*83 data is not shown due to its extremely low prevalence. Spheres in green indicate the presence of isolates with escape mutations for the allele family collected in that country. The sphere diameter is proportional to the total number of these isolates. Epitope escape substitutions and deletions for the eight allele family with fewest epitopes are listed on Supplementary Tables S6 and S7, respectively.

A small number of key viral changes may be sufficient to completely negate the contribution of these families, nearly devoid of SARS-CoV-2 ligands, to the cytotoxic protection. Substitutions and deletions (but no insertions) negated the binding of 42% and 37% epitopes (averaged by family), respectively, for weak alleles in the eight families with fewest alleles (Fig. 8B).

A pending issue is whether these SARS-CoV-2 isolates with changes that remove the HLA binding were collected from geographical zones with populations expressing these alleles. According to the Allele Frequency Database, the A*74, B*82 and C*18 families were prevalent in Africa whereas B*46 is common in Eastern Asia. These allele families were also common in minorities within these origins in other countries such as USA. By comparison, the B*52, B*73 and C*01 families were globally disseminated whereas B*83 was extremely rare. When isolates carrying mutations were geographically mapped using sample metadata and superimposed onto allele distribution, co-localization was observed in several cases (Fig. 8C). For instance, nine isolates from Ghana and USA showed the K369D (N protein) change, which canceled the binding of the 361-KTFPPTEPK-369 of eight A*74 alleles. Five isolates from Ghana and Kenya conveyed the ORF1ab deletion Δ6656-6744 (corresponding to Δ204-292 of nsp15_A1), which erased the 6669-AMDEFIERY-6677 epitope of the A*74:10 allele. Another example is constituted by nine isolates from India carrying the Q575R mutation in ORF1ab (Q395R in nsp2). This change invalidated binding to eight alleles of B*52 family, being India one of the countries with population samples enriched in this family. Likewise, the Δ6342-6432 deletion in ORF1ab (Δ417-507 in nsp14A2) was found in 17 isolates collected in Ghana and negated the 6353-TPAFDKSAF-6361 epitope of the B*82:03 allele. The deletion Δ872-966 (equivalent to Δ54-148 of nsp3) of ORF1ab underwent by two Hong-Kong isolates erased the 906-YLFDESGEF-914 epitope associated to three B*46 alleles. Finally, the M85Q (ORF1ab) substitution overrode five B*46 alleles and was found in Bahrain and USA isolates.

Another intriguing question is whether independent changes destroying two epitopes bound to alleles of any of these family tend to accumulate in the same isolates. Although it was a rare event, isolates with mutations negating two epitopes were identified in five out of the eight poor SARS-CoV-2-ligandome families. These isolates reached 2.59% of the total carrying at least one mutation negating B*52 epitopes. A prominent example is embodied by twenty-six isolates that combined alterations of ORF1ab Δ5828 (P504 in nsp13_ZBD protein) and large deletions in the ORF1ab 6341-6436 range (416-511 in nsp14A2_ExonN protein). These changes inactivated the 5827-NPAWRKAVF-5835 and 6353-TPAFDKSAF-6361 epitopes, respectively, of the B*82:03 allele. These isolates were collected from 22/03/20 to 09/02/21, in six USA states with different percentages of Afro-American population, suggesting some maintained dissemination degree and potential convergent selective pressure. The fact that these changes were also detected in isolation in several samples from the same country (22 and 62 isolates, respectively), indicates double mutants may have arisen by recombination.

## Discussion

This study aims to determine to what extend the mutations observed in large SARS-CoV-2 genome datasets can perturb the human cytotoxic response against this virus. This impact was studied in HLA class I molecules that practically cover the human population as a whole and, with special attention, to subsets with reduced SARS-CoV-2-ligand repertoires. In general, human and pathogen variability can greatly influence the CD8+ response, which may affect the outcome of infection. Some combinations of HLA class I haplotypes and viral genomes appear to further offset the balance towards an insufficient cytotoxic response and, thus, a probable bad prognosis. The surveillance of escape viral variants carried out in this study might therefore help to ameliorate enhanced susceptibility to COVID-19 in sub-populations by designing appropriate countermeasures.

The experimental evaluation of the immune response of every human allele associated to each viral variant is not feasible. Computational methods can facilitate this task and generate new, otherwise overlooked, hypotheses. Pioneering bioinformatic studies focused on predicting cytotoxic epitopes of a limited subset of common HLA alleles against the reference viral strain (13,20,23,26). However, SARS-CoV-2 has substantially evolved after more than a year of pandemic, resulting in a human-viral combination landscape of immense scale only approachable using automated techniques.

Bioinformatic approaches suffer from intrinsic limitations. These include the possible application of biologically inappropriate thresholds and potentially low predictive performance. Furthermore, alleles considered in algorithms as much as the priceless genome sampling by the worldwide sequencing effort still represent an underestimation of biological variability. Such obstacles were addressed in this study by: (i) utilizing an state-of-the-art algorithm that permits nearly universal fine-grained predictions (~3000 alleles); (ii) the application of stringent cutoffs that reflect the natural strictness of the ligand-HLA binding; (iii) the re-calculation of peptide binding affinity for each mutation; and (iv) the utilization of a large dataset of ~118,000 viral genomes and their corresponding metadata. Mutations were stratified by occurrence, reduction of HLA-binding affinity and geographical dissemination. Thus, the integration of omic data and immunoinformatics in this study very likely capture, despite drawbacks, the principal trends that respond to the posed questions.

Large epitope numbers were computationally predicted to be presented by most supertypes. Although all these supermotifs appeared mutated in at least one isolate, most of these mutations did not overcome the supermotif degeneracy. In most cases, the HLA binding affinity was reasonably maintained except from (i) residue substitutions in the second and C-terminal positions of the ligand core, amino acids that usually are anchor motifs; and (2) large deletions that fully removed the epitope. For instance, the Spike-W152C mutation and deletions in the 6342-6432 range in ORF1ab removed several epitopes at the level of supermotifs, and were coupled to several other changes. Respect to the persistence of these escape mutations, point substitutions are likely less prone to impose a dramatic fitness although some extensive deletions have been also been shown to be compatible with infection and transmission (27,28). Large deletions have been related to progressive adaptation to host and reduced virulence (29,30), but their middle-term stability should be analyzed case-to-case.

A central question is whether escape mutations have longitudinally accumulated in genomes of individual isolates. If so, such emerging strains would have acquired, or be in the process of acquiring, enhanced capacity to infect individuals previously able to mount an effective cytotoxic response. However, the emergence of this challenging phenotype would not be expected after the examination of the genomic space of the virus carried out in this study. Even the forward line of mutated variants in this respect only combined low numbers (<15%) of escape supermotifs of a given supertype. The remaining intact supermotifs, other HLA class I loci and heterozygosity should compensate escape mutations, provided that the pool of naïve lymphocyte is high enough and the innate-to-adaptive response priming correctly coordinated. Notably, the humoral and CD4^+^ responses would likely remain active and be sufficient in many cases. Therefore, we conclude that the systemic nature of the immune response translates into most healthy subjects remaining competent to respond against variants. The only exception that moderately threatens supertype redundancy was the B27 supertype with isolates that convey evading mutations for up to 33% of these supermotifs. This supertype is common in many populations such, in particular, in Eskimo (31), which may be exposed to “Northern America” isolates with disabled B27 supermotifs.

The emergence of isolates that undergo the step-wise accumulation of genetic markers to achieve extended cytotoxic resistance should not be ruled out. This may be favored by considering the explosive expansion of the virus worldwide. However, the mutational space would be reduced in practice due to potential antagonism between cytotoxic evasion pressure and structural-functional restrictions of proteins. However, a sizable fraction of the human population has been infected with the virus, which represents innumerable replication cycles and infection attempts. Some variants have been linked by other scientific groups to different clinical phenotypes such as increased mortality (32) and antibody escape (33). Likewise, progressive mutation and recombination in SARS-CoV-2 may conceivably achieve a critical number of supermotif escape mutations that collectively constitute a selective advantage. Some identified isolates appeared to have experienced a higher-than-expected number of these changes over the genetic noise, and may have initiated the evasion-driven process.

On the other hand, according to our computational study, a worrisome scenario has already occurred for around ~10% of alleles able to bind a reduced number of ligands from the SARS-CoV-2 proteome. Among them, the A*74, B*82 and C*18 allele families, with sub-Saharan African origin, and the C*46 family, with Far East origin, excelled. Lost or debilitation of the cytotoxic response would make these individuals too dependent upon the humoral response, which can be inefficient during primary infection in some cases (1). This may be very relevant when these alleles are combined into the same haplotype, in particular when in homozygosis.

Underprotection may become exacerbated if these individuals become infected with these escape variants. Given their low epitope redundancy, a very few number of viral mutations, such as those identified in this study, may suffice to circumvent both the cytotoxic primary and memory T responses. The geographical co-existence of viral variants that experience epitope switch respect to some HLA class I molecules and individuals expressing these alleles may exert immediate selective pressure. This may cause rampant dissemination of emergent strains in these niches with local clinical consequences. Most isolates at great risk of achieving critical mutations to impair the CD8^+^ ligand repertoires in these families were found in “Northern America” where some African Americans and Asian subpopulations carried these alleles. Whether these immunotypes with further diminished SARS-CoV-2 ligandomes have undergone positive selection warrants massive local HLA genotyping and viral sequencing. Some of these alleles may be ancestrally specialized in single pathogens, but unable to be effective against international viral infections as reported for Dengue (34), HIV (35) and influenza (36).

In conclusion, here we provide a complete repository of the predicted escape mutations in a recent NCBI genome sampling of SARS-CoV-2. Fortunately, accumulation of these mutations in single isolates does not appear close enough yet to be alarming at the global population level. However, isolates carrying mutations able to override limited CD8^+^ response in some alleles and haplotypes are already co-circulating with individuals carrying these HLA class I molecules. Emerging SARS-CoV-2 variants may further increment the susceptibility of highly vulnerable communities and should be actively surveyed to coordinate appropriate countermeasures. In this respect, bioinformatic pipelines operating on a timely basis may play an irreplaceable role in the protection against this and other pandemic threats.

## Materials and Methods

### Data acquisition

SARS-CoV-2 coding sequences and isolate metadata were downloaded from the NCBI repository (Last accession: 19/03/2021) (37). Protein sequences of clinical isolates showing length differences >3% respect to the reference variant were considered anomalous and rejected. The country of origin of isolates were assigned to sub-continental regions following the M49 United Nation geoscheme. Experimental epitopes were downloaded from the IEDB (Last accession: 19/03/2021)(16) using the following search terms: Epitopes: “Any epitopes”; Assay: “T Cell”, “MHC Ligand” and outcome: “Positive”; MHC Restriction: “MHC Class I”; Host: “Human”; Disease “COVID-19 (ID: DOID: 0080600)”.

Alleles for the twelve HLA class I supertypes were acquired from the original publication (12).

Geographical localization of populations with allele families with few epitopes was carried out using the Allele Frequency Net Database (38). Only samples with at least 50 individuals and ≥1% frequency for the given allele family were considered.

### HLA class I epitope prediction and analysis

HLA class I epitopes between 8-12 residues in 11 viral proteins, and the ORF1ab polyprotein, of the SARS-CoV-2 reference proteome (Wuhan-1; RefSeq: NC_045512.2) were predicted for 2,915 alleles using NetMHCIpan EL 4.1 (22). Binding epitopes were considered those that satisfy the rank ≤ 0.5 (EL rank) and score (EL score) ≥ 0.5 estimations provided by this neural network method. The predictive performance of this algorithm was superior when trained with mass spectrometry elution (EL) data than when trained with binding affinity (BA) data and therefore the former is recommended by the developers for general applications. However, the “EL score” quantifies biologically meaningless abstract units whereas the score of the BA version “BA score” reflects the IC50 in nM. Thus, to take advantage of the strengths of both strategies, the approximate equivalences between EL and BA scores were assessed by exponential regression (r^2^=0.69) (Supplementary Fig. S1). This comparison resulted in a value for “EL score” ≥ 0.5 was roughly equivalent to an IC50 ≤ 500nM. This affinity threshold is satisfied by most medium to high-affinity real ligands (39). Redundant epitopes with distinct lengths and lower “EL scores” but sharing the same peptide core and allele were ignored.

Intra-family coherence was calculated by comparing the non-redundant epitope pools between all alleles of the same family and calculate the average Jaccard coefficient (intersection divided by the union) of all families. For that, the intersection was constituted by the number of epitopes that showed a perfect coordinate match for two alleles among all epitopes identified by each allele. Inter-family epitope correlation was calculated by comparing family epitope pools, i.e. those shared by at least half of the alleles of the alleles in the family, like explained above between families. A matrix with all inter-family Jaccard coefficients was used for agglomerative hierarchical clustering by *clustermap* function of *seaborn* data visualization Python library with default options.

Supermotifs, or supertype-associated epitopes, were those predicted as non-redundant epitopes showing perfect coordinates for ≥ 50% alleles in the supertype (12). Only alleles which motifs were experimentally established or shared exact match(es) to second and C-terminal peptide positions, i.e. B and F pockets of the HLA class I groove, in the original reference were considered.

### Mutation analyses

Mutations respect to the proteins of the reference Wuhan-1 strain (RefSeq: NC_045512.2) were identified by aligning with Clustal Omega 1.2.1 (40) with an in-house perl script. All adjacent insertion or deletion runs were collapsed into single events. Non-conservative mutations were deemed those involving distinct physicochemical classes: acidic (D, E), amide (N, Q), basic (H, K, R), cysteine (C), glycine (G), hydroxyl (S, T), hydrophobic aliphatic (A, I, L, M, V), hydrophobic aromatic (F, W, Y) and proline (P) residues.

The impact of point substitutions on epitope binding was assessed by recalculating the “EL score” and “El rank” of the mutated peptide. For insertions, flanking regions to the insertion limits were taken to complete 22mer sections and binding also recalculated. Likewise, for deletions, the resulting 11mers flanking the deletion limits were merged into 22mer sections. Based on the “BA score” and “EL score” correspondences, “EL scores” of < 0.1 roughly corresponded to BA scores of >5000nM (Supplementary Fig. S1), associated to non-binders according to the IEDB curators. Thus, mutations causing medium-strong peptide binders decreases to EL scores < 0.1 besides EL rank ≥ 1 were deemed epitope escape mutations. For supermotif escape mutations, the allele of the supermotif showing the highest “EL score” for the wild-type epitope was tested. For the eight families with fewest epitopes, escape mutations were calculated for each allele.

Network graphs of coupled mutations were carried out using the NetworkX (41) Python library.

## Author Contributions

**Conceptualization:** Daniel López, Michael J. McConnell, Antonio J. Martín-Galiano.

**Data curation:** Anna Foix, Antonio J. Martín-Galiano.

**Formal analysis:** Anna Foix, Antonio J. Martín-Galiano.

**Funding acquisition:** Daniel López, Michael J. McConnell, Antonio J. Martín-Galiano.

**Investigation:** Anna Foix, Daniel López, Michael J. McConnell, Antonio J. Martín-Galiano.

**Methodology:** Anna Foix, Antonio J. Martín-Galiano.

**Project administration:** Antonio J. Martín-Galiano.

**Resources:** Antonio J. Martín-Galiano.

**Software:** Anna Foix, Antonio J. Martín-Galiano.

**Supervision:** Daniel López, Michael J. McConnell, Antonio J. Martín-Galiano.

**Validation:** Antonio J. Martín-Galiano.

**Visualization:** Anna Foix, Antonio J. Martín-Galiano.

**Writing–original draft:** Antonio J. Martín-Galiano.

**Writing–review & editing:** Daniel López, Michael J. McConnell, Antonio J. Martín-Galiano.

## SUPPORTING INFORMATION CAPTIONS

Figure S1. Correspondence between netMHCpan 4.1 EL and BA scores.

Table S1. Intra-family conserved epitopes.

Table S2. Supertype-associated epitopes.

Table S3. Supermotif escape substitutions.

Table S4. Supermotif escape deletions.

Table S5. Isolates carrying escape mutations for five or more supermotifs.

Table S6. Escape substitutions for allele families with few epitopes.

Table S7. Escape deletions for allele families with few epitopes.

