## Supplementary Figure S1 for "Predicted Impact of the Viral Mutational Landscape on the Cytotoxic Response against SARS-CoV-2"

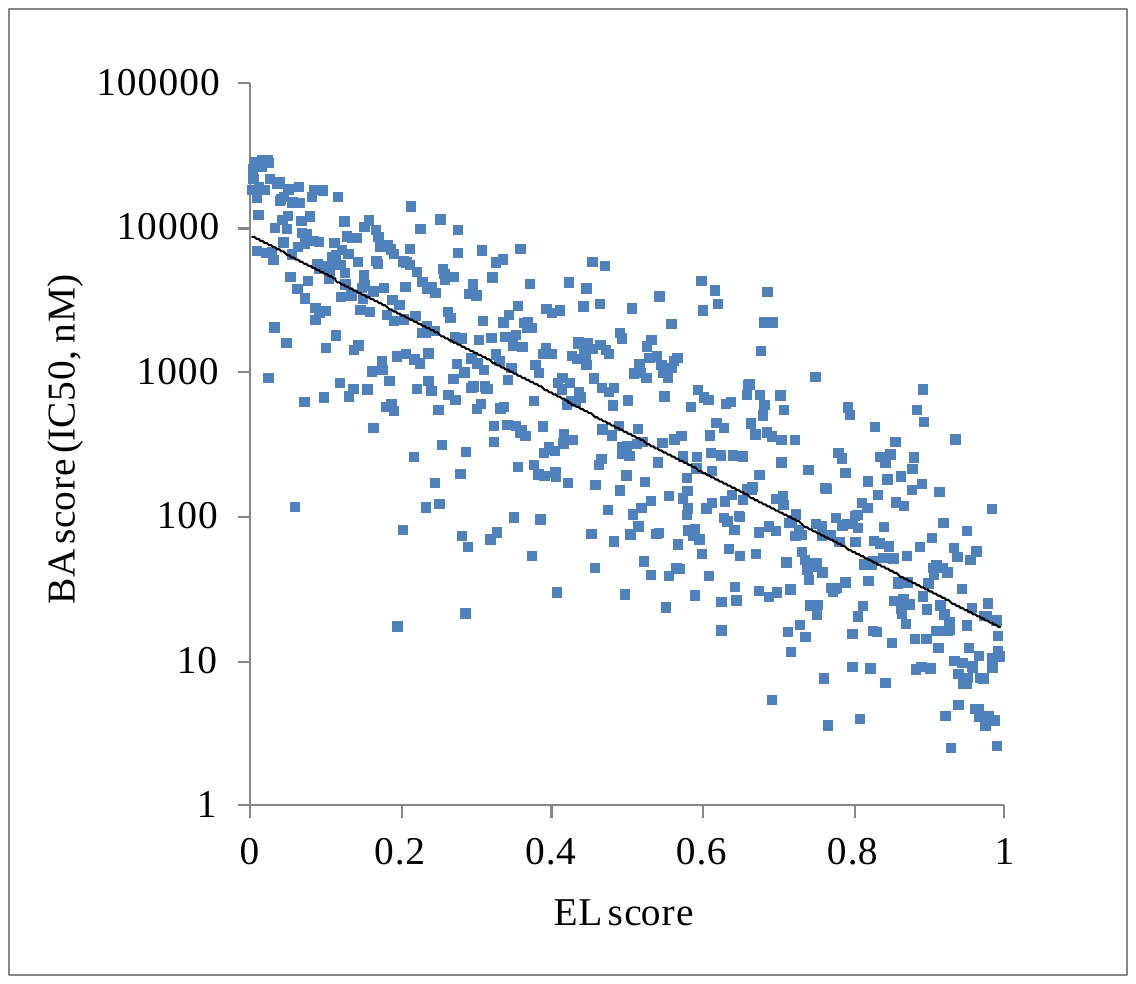

**Figure S1. Correspondence between netMHCpan 4.1 EL and BA scores**. Three random SARS-CoV-2 HLA class I nonamers predicted by netMHCpan 4.1 EL for each 0.005 score interval between 0 and 1 were selected. Binding of the same nonamer dataset was re-calculated for the same allele using netMHCpan 4.1 BA.
